# Tradeoffs in planning marine protected areas for kelp forest resilience: protecting climate refugia is not always the best solution

**DOI:** 10.64898/2026.04.01.715997

**Authors:** Jess K. Hopf, Anita Giraldo-Ospina, Jennifer E. Caselle, Kristy J. Kroeker, Mark H. Carr, Alan Hastings, J. Wilson White

## Abstract

Marine protected areas (MPAs) are increasingly promoted as climate mitigation tools, yet guidance on their placement to maximize resilience against climate stressors like marine heatwaves remains limited. Here, we develop MPA placement guidelines that explicitly consider a mechanistic pathway through which MPAs could enhance kelp forest resilience to heatwaves: protecting fishery-targeted urchin predators to prevent kelp overgrazing. Using a spatially explicit, tri-trophic model of California kelp forests, we evaluate alternative MPA configurations across a hypothetical coastline where half the habitat experiences an increased probability of experiencing heatwaves. We found that effective MPA placement depends on whether MPAs are being newly established or reconfigured within an existing network, and that among-patch connectivity and spillover played vital roles in the relative effectiveness of different MPA configurations. Changes in resilience occurred primarily at the patch scale, with trade-offs between increased within-MPA resilience and decreased resilience in some fished areas, resulting in minimal coastwide population effects. For example, for new MPAs, large single MPAs within heatwave-prone areas maximized within-MPA resilience gains, while multiple small MPAs in heatwave refugia best supported whole-coast resilience. When reconfiguring established networks, expanding existing MPAs in refugia areas was most effective. We also demonstrate the importance of considering MPA recovery timescales: for example, relocating old MPAs to heatwave refugia yielded minimal short-term benefits due to the loss of rebuilt, previously fished, predator biomass. Our findings demonstrate that climate-adaptive marine planning should explicitly consider the spatiotemporal implications of trophic cascades, connectivity, and transient population dynamics to support ecosystem resilience.

## Introduction

The accelerating pace of climate change underscores the urgent need to identify local actions that may mitigate its impacts. A key challenge is the increasing frequency, duration, and severity of marine heatwaves (hereafter ‘heatwave(s)’), which pose significant risks to marine ecosystems and the services they provide (Frölicher et al., 2018; K. E. Smith et al., 2021, 2023). Ultimately, the remedies for climate change are global, but there is also a pressing need for local mitigation strategies. For example, there is currently much interest in the role marine protected areas (MPAs) may play in mitigating climate impacts like heatwaves (Jacquemont et al., 2022; Roberts et al., 2017; J. G. Smith et al., 2023; J. W. White et al., 2025), even though MPAs have historically been established to protect marine ecosystems from different local stressors, such as fishing (Lubchenco & Palumbi, 2003).

A plausible mechanism through which MPAs – especially no-take MPAs, which is our focus – may mitigate heatwave impacts is by rebuilding and protecting higher-trophic species targeted by fisheries, leading to increased ecosystem resistance and resilience through trophic cascades (Jacquemont et al., 2022; Kumagai et al., 2024; J. W. White et al., 2025). For example, in many kelp forests, abundant predators can keep populations of herbivorous sea urchins in check, reducing the likelihood of shifting from a productive kelp-dominated state to a less desirable, overgrazed urchin barren (Eisaguirre et al., 2020; Hamilton et al., 2023). Heatwaves can affect this dynamic by reducing kelp growth and recruitment (Hollarsmith et al., 2020; Michaud et al., 2022; Zimmerman & Kremer, 1986), leading to a reduced production of detached kelp fronds (’drift kelp’), a preferred food source for urchins. This loss of drift kelp, combined with heatwave-induced increased grazing rates, can trigger urchins to switch from feeding cryptically on drift in rocky crevices to roaming exposed on the reef, targeting standing kelp stipes (Kriegisch et al., 2019; Rennick et al., 2022; J. G. Smith & Tinker, 2022) (Fig 1). Because kelp forest resilience to heatwaves is a fine balance of consumer-resource dynamics, using MPAs to protect and rebuild fishery-targeted urchin predator populations may buffer against heatwave-induced kelp loss (Hopf et al., 2025). This benefit of MPAs was documented empirically in California, USA, where MPAs that protected urchin predators were less likely than fished areas to lose kelp biomass during a heatwave (Eisaguirre et al., 2020; Kumagai et al., 2024).

**Figure 1:**
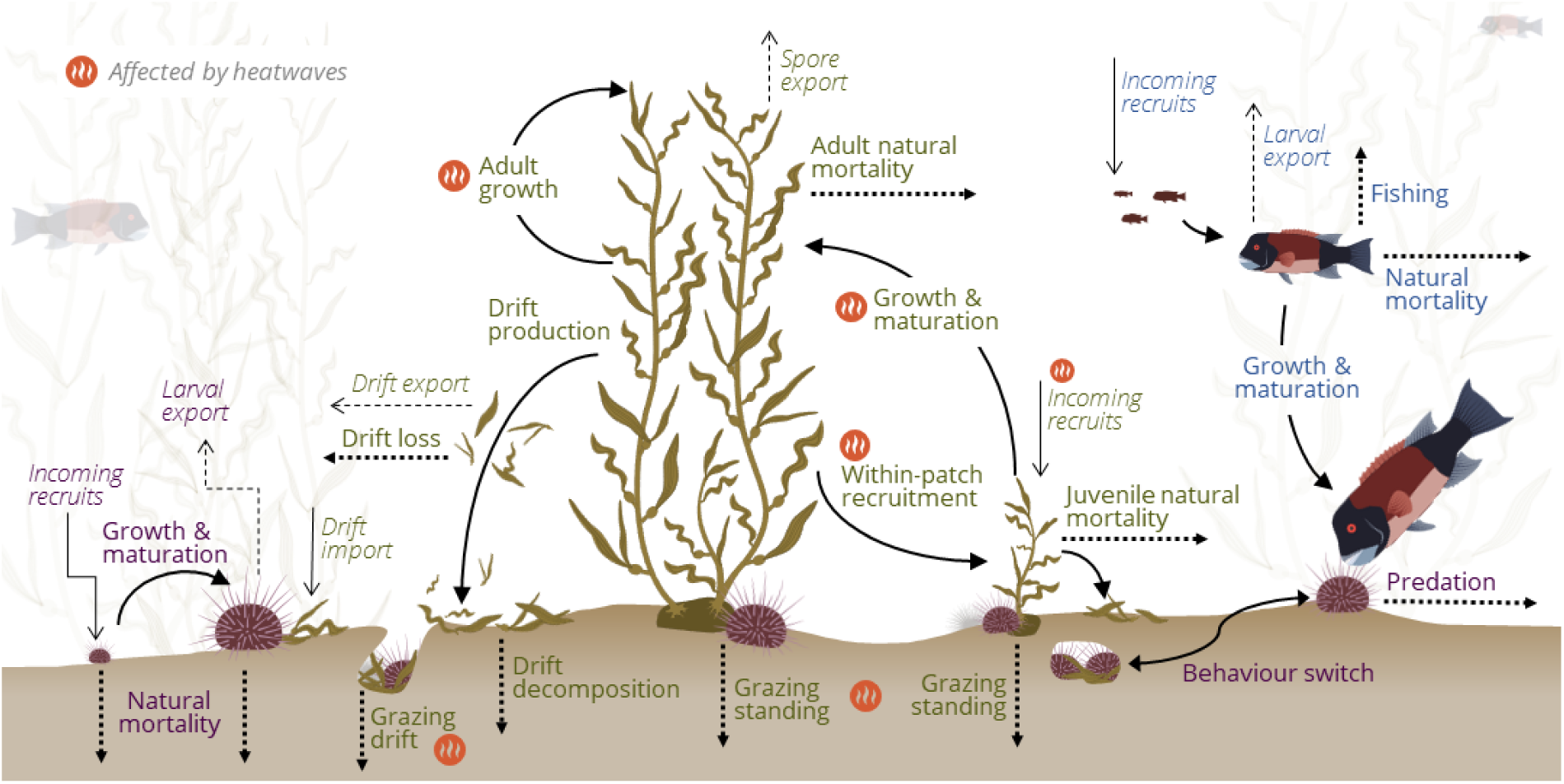
Three-species model overview. Key patch-level processes, including those affected by heatwaves (orange symbols), captured in our kelp-urchin-predator population model. Purple and green text show processes relevant to the kelp-urchin sub-model, and blue text indicates processes relevant to the predator sub-model. Dashed lines indicate export/loss, and solid lines indicate import/growth process. Non-italicized and italicized text indicate within- and among-patch processes, respectively. All artwork by Jess K. Hopf, except giant kelp icons, which are modified from artwork by Jane Thomas sourced from ian.umces.edu/media-library under the CC BY-SA 4.0 license.

As MPA networks have typically been designed using biophysical guidelines (e.g., habitat representation, connectivity) that assume stationary climate conditions, climate change impacts and the mechanisms of climate resilience have not been explicitly considered in traditional MPA planning (Lopazanski et al., 2023). In contrast, emerging climate-focused conservation planning (often termed “climate-smart” or climate-adaptive planning) aims to incorporate projected spatial and temporal patterns of climate change into MPA design (e.g., Arafeh-Dalmau et al., 2023; Brito-Morales et al., 2022; Buenafe et al., 2025). These approaches typically prioritize areas of reduced climate exposure, climate refugia, or high connectivity under future scenarios, and are increasingly promoted in international policy, including the post-2020 Global Biodiversity Framework (COP15 2022). However, while climate-smart planning could provide important guidance on where MPAs might best be placed under climate change, this approach has generally focused on metrics of exposure to perceived climate stressors (e.g., changes in the mean or variance of sea surface temperature; Brito-Morales et al., 2022) rather than explicitly considering the ecological mechanisms that may promote resilience to specific climate stressors such as heatwaves (but see White et al., 2025).

Here, we develop MPA placement guidelines for future marine heatwaves that explicitly consider the mechanistic pathway through which MPAs may enhance resilience to heatwave-driven kelp forest collapse by protecting fishery-targeted urchin predators. We focus on the near-term (20-year) consequences of alternative spatial designs for new and established MPA networks, accounting for transient dynamics following MPA establishment. Because the biomass and abundance of previously fished predator populations typically recover over years to decades (Hopf et al., 2016; J. W. White et al., 2013), resilience benefits arising through trophic pathways may also be delayed. These lags are particularly important when 1) setting expectations for the assessment of MPA resilience outcomes, and 2) considering change to MPA network design (CDFW, 2022). In the case of MPA relocation, there may be trade-offs between reduced future climate exposure (if, for example, shifting to a climate refugia), the time required to rebuild predator populations in the new location, and the loss of conservation benefits from opening established MPAs to fishing. Specifically, we ask, where should new MPAs be placed relative to regions of higher/lower heatwave probability, as well as other existing MPAs, to maximize kelp forest resilience?

To answer this question, we use a three-species, multi-patch (along an idealized linear coastline) population model that captures the key species and trophic interactions in southern California kelp forests: giant kelp (*Macrocystis pyrifera*), herbivorous purple urchins (*Strongylocentrotus purpuratus*), and a fishery-targeted predatory fish (California Sheephead; *Bodianus pulcher*, henceforth “Sheephead”). This is a spatial extension of a model previously used to investigate single-population heatwave impacts (Hopf et al., 2025). Here, we compared the short-term resilience consequences of alternative no-take MPA placements in response to an increased incidence of heatwaves. We consider multiple spatial configurations, including scenarios without previously established MPAs in the region and scenarios with an established MPA network comprised of two individual MPA patches covering 12.5% of the kelp-forest habitat (similar to the situation in California, Fig 2). We considered a scenario in which half of the coastline has an increased chance of experiencing a heatwave (25% chance each year), while the other half is a refugium habitat that experiences historical levels of environmental variability.

**Figure 2:**
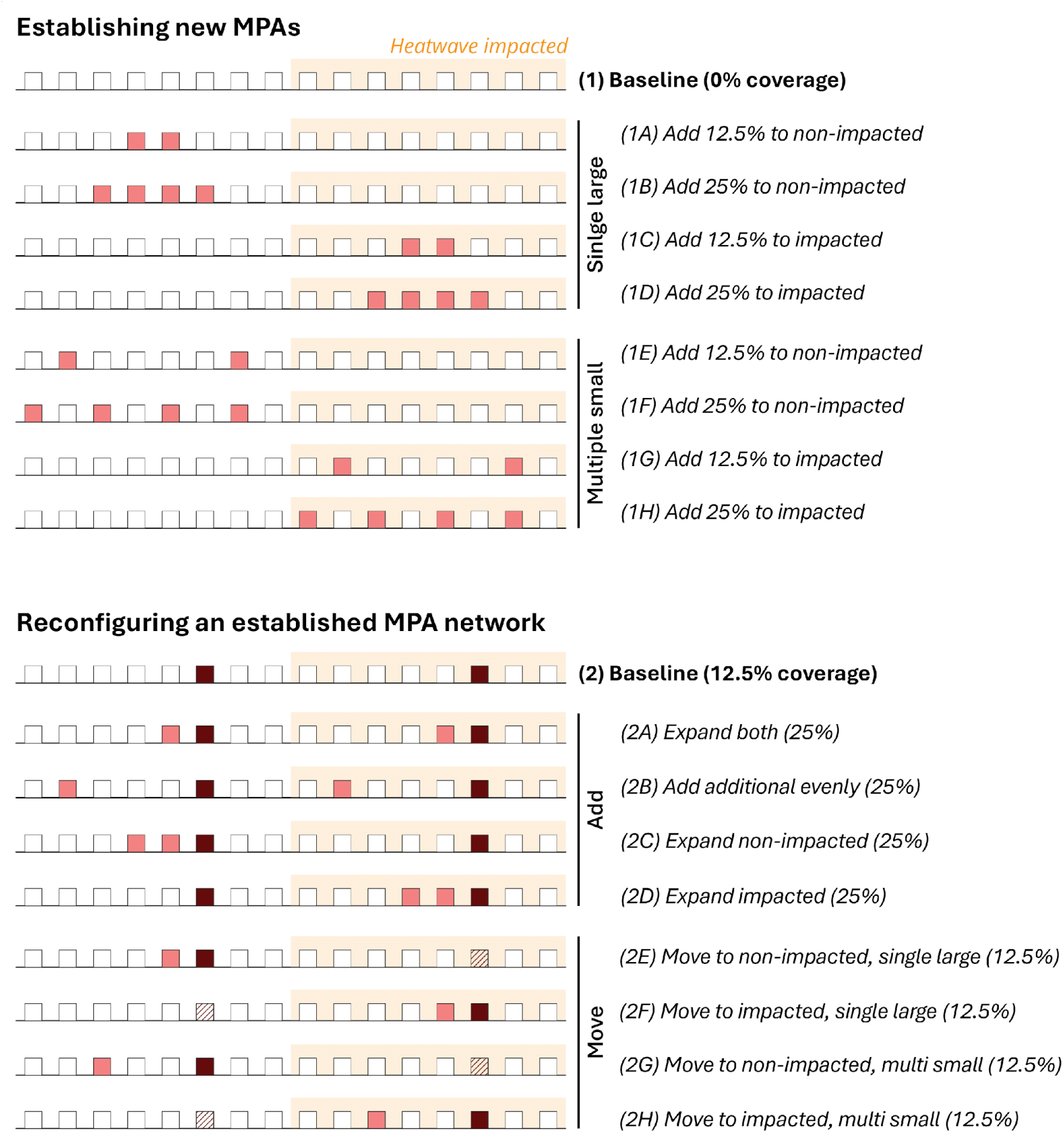
MPA scenarios. Schematic of the modelled, linear coastline with the MPA scenarios considered in this study, including establishing new MPAs (light red squares) along a coastline without any existing MPA network, and adding or relocating MPA patches in a network with established MPAs (dark red and hatched squares). Each square represents a single kelp forest patch covering 6.25% of the total kelp forest area. Patches are separated by sandy substrate along a infinite coastline (so the leftmost and rightmost patches depicted here are also neighbors). Orange regions indicate patches impacted by marine heatwaves, while the non-highlighted region is a heatwave refugium.

## Results

We quantified the ‘resilience’ of a given kelp forest habitat patch as the proportion of time kelp biomass was above a minimum threshold (1% of no-heatwave, kelp-forested state) during the first 20 years after the MPA configuration was changed (and the heatwave probability had increased for half of the coastline). This definition encompasses both the resistance of kelp-forest patches to a state change and the recovery of degraded patches to a kelp-dominated state (primarily through kelp spore and drift imports). As our model includes natural variability in larval recruitment (and the incidence of heatwaves; see Methods), there is a chance of kelp forest decline independent of heatwave perturbation. Therefore, our definition of resilience includes overall resilience to change, not just resilience to disturbances. Distributions of resilience across the 5,000 simulation runs were bimodal, reflecting phase-shifting system behavior between stable kelp-dominated forests and persistent urchin barrens (SI Figs). To address our research questions, we focus on the absolute difference between the median resilience values of the baseline and alternative MPA configurations, on both patch-by-patch and whole-coastline scales (Figs 3-4).

**Figure 3:**
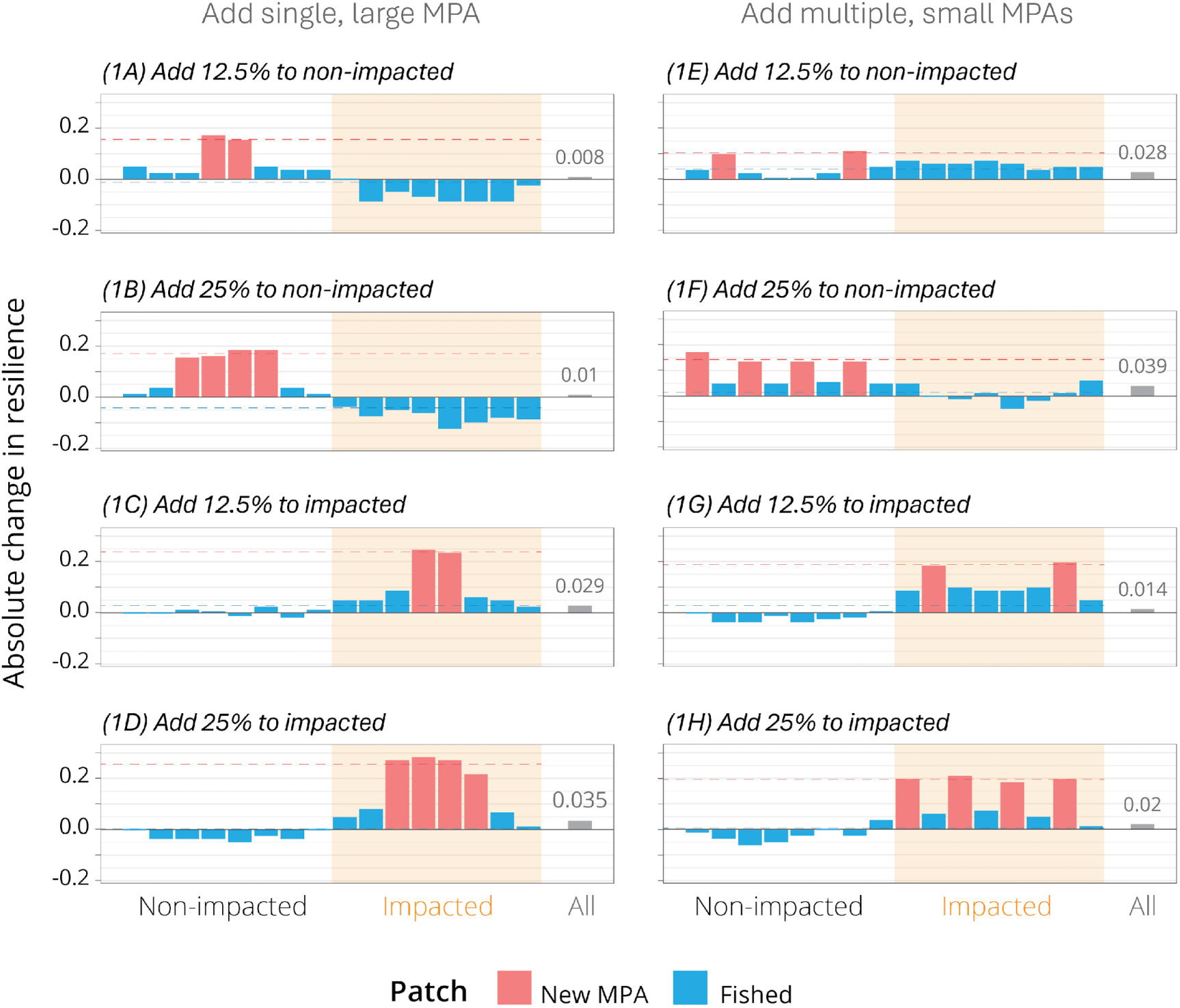
Resilience changes when establishing new MPAs. The absolute change in median resilience (proportion of years kelp-forest patches spent above the kelp biomass threshold, which is 1% of the no-heatwave, kelp-forested state), 20 years post-MPA implementation. Values are relative to the baseline scenario of no MPAs. Each blue/red bar is an individual patch (6.25% of the total kelp forest area), ordered along an infinite modelled coastline that is partially affected by marine heatwaves (orange box indicates impacted patches). Grey bars and text are changes in median resilience at the population scale. Horizontal dashed lines represent mean changes in patch resilience for MPA (red) and fished (blue) patches (i.e., means of bar heights). Letters relate to scenarios outlined in Figure 2. See also Supplementary figure x.

**Figure 4:**
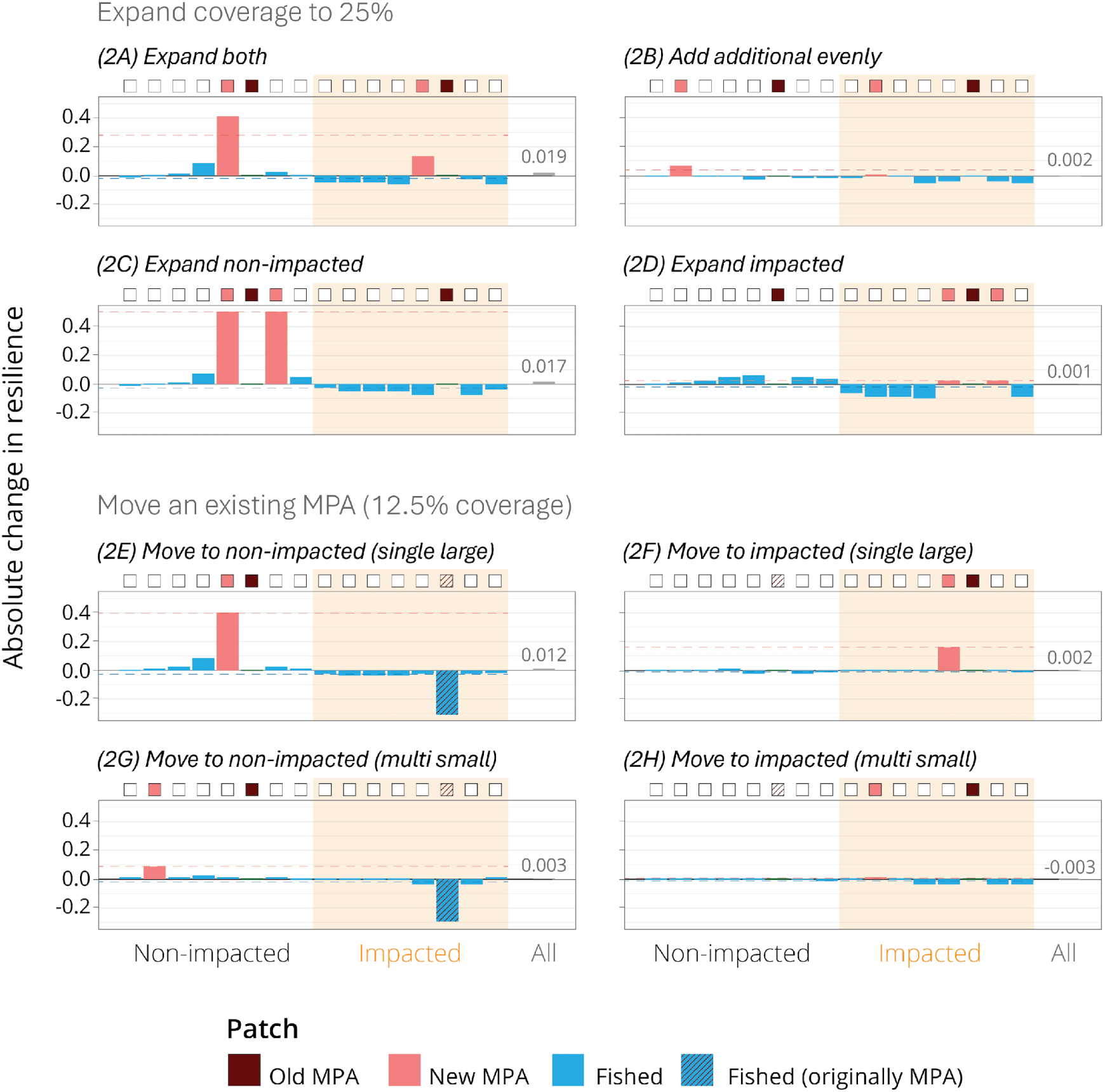
Resilience effects of reconfiguring an established MPA network. The absolute change in median resilience (proportion of years kelp-forest patches spent above the kelp biomass threshold, which is 1% of the no-heatwave, kelp-forested state), 20 years post-MPA implementation. Values are relative to the baseline scenario of two old MPAs. Each blue/red bar is an individual patch (6.25% of the total kelp forest area), ordered along an infinite modelled coastline that is partially affected by marine heatwaves (orange box indicates impacted patches). Grey bars and text are changes in median resilience at the population scale. Horizontal dashed lines represent mean changes in patch resilience for MPA (red) and fished (blue) patches (i.e., means of bar heights). Letters relate to scenarios outlined in Figure 2. See also Supplementary figure x.

Habitat patches with newly placed MPAs had varied resilience gains depending on MPA coverage, configuration (single large or several small), and placement, as well as adjacency to an existing MPA patch or high heatwave likelihood (red bars in Figs 3-4). Notably, these gains within the MPA came at a cost to resilience in some, but not all, fished patches (blue bars in Figs 3-4). Typically, fished patches closer to new MPA patches experienced resilience gains, while more distant fished patches had resilience losses. This is because there is spillover of kelp spores and drift kelp from healthy kelp forests inside MPAs to neighboring patches, but even redistribution of fishing pressure from new MPAs into the remaining fished habitat. These trade-offs among patches resulted in none to minimal (<0.05 unit change) increases in median resilience for the whole population after adding/moving MPAs (grey bars in Figs 3-4). Within-patch and population effects were consistently greater with larger MPA coverage (Figs 3-4).

Single large MPAs resulted in consistently greater resilience gains within MPA patches than multiple small MPA patches of the same coverage and placement relative to the heatwave-impacted area. This was true for both establishing a new MPA network (compare red horizontal dashed lines across columns, Fig 3) and reconfiguring an existing one (e.g., compare red horizontal dashed lines between scenarios 2A and 2B or 2E and 2G in Fig 4). However, single large MPAs also resulted in stable or declining resilience in fished patches, on average. For example, no fished patches had reduced resilience when establishing two separate, single-patch MPAs (covering 12.5% of the total area) in the non-impacted, heatwave refugium area (1E in Fig 3), but over half the fished patches saw declines in resilience when establishing a single large MPA covering 12.5% in the heatwave refugium (1E in Fig 3). This difference arose because spillover of kelp forest benefits is shared between habitat patches in the same large MPA but not between distant small MPAs.

When considering placement relative to the heatwave-impacted region, we found that establishing new MPAs (without existing MPAs) in the heatwave-impacted region resulted in greater within-MPA gains than establishing them in the non-impacted refugium area, regardless of configuration (e.g., compare 1A, 1C, 1E, and 1G in Fig 3). Conversely, in the scenario with existing MPAs, adding new MPAs or moving existing ones to the heatwave refugium area resulted in markedly greater resilience gains than focusing MPA patches in the impacted region (e.g., compare 2C to 2D, and 2E to 2F in Fig 4).

The effects on fished-patch resilience of MPA placement relative to the heatwave region interacted with MPA configuration: resilience losses occurred and were greatest with single large MPAs in refugia areas, regardless of whether there were previously established MPAs (1A, 1B in Fig 3, and 2A, 2E in Fig 4). Critically, heatwave-impacted patches that were originally MPAs but then opened to fishing saw the greatest decreases in resilience (2E, 2G in Fig 4), while refugia MPA-to-fished patches saw no change (2F, 2H in Fig 4).

Gains in median resilience at the population scale were likewise influenced by an interaction between MPA configuration (single large, or multiple small) and placement relative to the heatwave-impacted region (grey bars Figs 3-4). When establishing new MPAs, the greatest increases in resilience occurred with larger coverages (25%) either with a single MPA in the impacted area, or multiple MPAs in the non-impacted area (1F, 1D in Fig 3). When reconfiguring or adding new MPAs to the established MPA network, we found that expanding existing MPAs (adding new or moving old MPA patches) had the largest resilience gains across the whole population, but only when at least half of the final coverage was in the heatwave refugium (2A, 2C, 2E in Fig 4). Moving established MPAs had little benefit (2E-2H in Fig 4).

## Discussion

MPAs can enhance resilience against heatwave-driven kelp forest collapse by protecting and rebuilding fishery-targeted urchin predators (Eisaguirre et al., 2020; Jacquemont et al., 2022; Kumagai et al., 2024). By explicitly modelling this mechanistic pathway and evaluating alternative MPA configurations, we show that the extent of the climate mitigation benefit that could be expected depends not only on where MPAs are placed relative to climate refugia but also on spillover^1^ and connectivity processes, transient dynamics, and existing MPA protection. Critically, resilience responses were highly spatially heterogeneous and emerged primarily at the patch level, rather than the entire population, with trade-offs between increased resilience within-MPAs and decreased resilience in some (but not all) fished patches leading to minimal overall changes in resilience at the whole-coast scale. This lack of whole-coast resilience benefits from MPAs reflects current understanding of population dynamic responses to MPAs in systems without severe overfishing: there should be local within-MPA increases in biomass but not substantial whole-population biomass increases (e.g., Ovando et al., 2021; White et al., 2025).

The management implications of our findings will depend on planning objectives; for example, if a climate-adaptation goal of a new MPA network is to maximize resilience against heatwave-driven kelp forest collapse primarily *within* MPA areas, then establishing a large single MPA in heatwave-vulnerable areas may be best (Fig 5). In the fished area, however, this approach will have minimal benefits, which will diminish further with larger MPA coverages. Conversely, if the goal is to use MPAs to support more uniform resilience across the whole coastline, then multiple small MPAs in the heatwave refugia area (or less so, the impacted area) may be more suitable (Fig 5). This work provides starting foundations for such considerations to be folded into climate-adaptive planning alongside climate-refugia approaches (e.g., Brito-Morales et al., 2022), risk-spreading strategies (e.g., Allison et al., 2003), dynamic MPAs (e.g., Tittensor et al., 2019), and other mitigation strategies (Arafeh-Dalmau et al., 2023; Buenafe et al., 2025).

**Figure 5:**
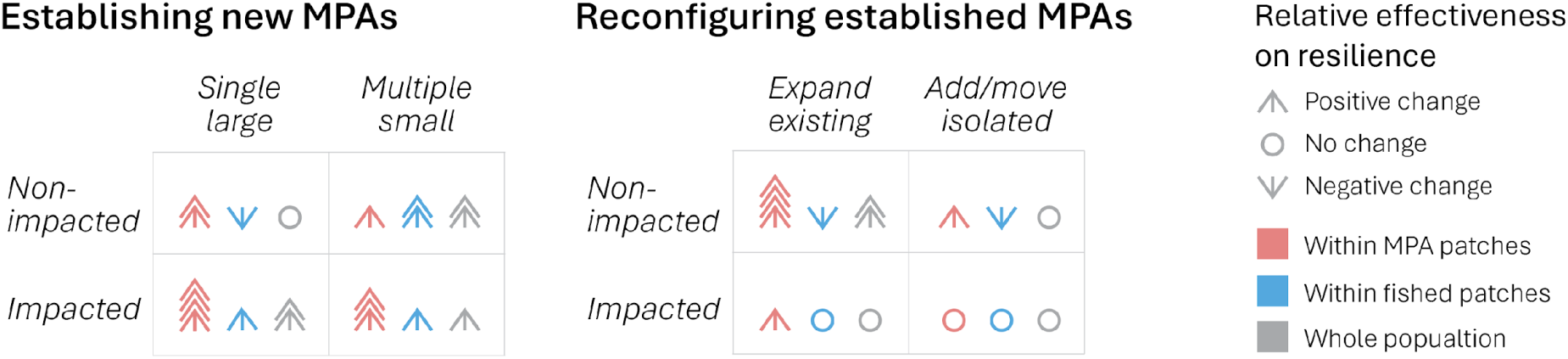
Summary of guidelines for resilient kelp-forest MPAs. Overarching guidelines for establishing new MPA networks and reconfiguring (expanding or relocating patches) existing MPA networks based on changes to resilience (proportion of years kelp-forest patches spent above the kelp biomass threshold) within kelp-forest patches and across a linear coastline. ‘Non-impacted’ and ‘impacted’ refer to where the newly established MPA patches are focused relative to heatwaves. Symbols qualitatively indicated the effectiveness of a given strategy relative to the baseline and other scenarios, with larger arrows indicating greater relative change.

Among-patch connectivity and spillover played a vital role in the relative effectiveness of different MPA configurations in our analysis. Whether changes in the MPA configuration increased or decreased the resilience of fished patches reflected a balance between exported benefits from MPAs (increased kelp spore supply and drift kelp biomass, which primarily affected neighboring patches) and the even redistribution of fishing pressure from newly closed areas to the remaining open patches. As such, fished patches adjacent to newly established and long-standing MPAs typically exhibited increased resilience, whereas patches further away often experienced decreases. To our knowledge, this is the first study to explicitly call attention to potential negative resilience outcomes associated with MPA configurations under climate stress. Such outcomes are consistent with the expected implications of MPAs on fisheries (Botsford et al., 2009; White et al., 2014), in which redistributed fishing pressure can temporarily or permanently reduce expected fishery yields. The spillover of MPA benefits (kelp spores and larvae) between patches is also a primary driver behind our finding that expanding established MPAs was notably more effective at increasing resilience within those MPAs, than placing distant MPAs.

Our results demonstrate the importance of accounting for ecological recovery timescales when evaluating the climate-mitigation potential of MPAs. The mechanistic pathway for resilience we focused on requires sufficient biomass of fished predators to reduce urchin numbers below thresholds (Ling et al., 2015). While the time required to reach these biomasses is unknown (and will depend on pre-protection fishing levels; Nickols et al., 2019), MPA literature indicates that it may take years to decades to rebuild previously fished populations to pre-exploitation levels (Claudet et al., 2008; Hopf et al., 2016; J. W. White et al., 2013). Indeed, older MPAs have demonstrated greater resilience to kelp loss than younger ones (Eisaguirre et al., 2020). In our model, we assume that old MPAs had fully rebuilt Sheephead biomass and population structure before increasing heatwave frequency. Under these conditions, we found that offsetting the cost of relocating these old MPAs (loss of established biomass) may occur only in limited situations: specifically, if there is an existing MPA in the non-impacted area that can be expanded to maintain overall coverage (scenario 2E in Fig 4). Notably, we found that moving an established MPA to a refugia area, but distant from other established MPAs, was unlikely to have notable benefits (scenario 2F in Fig 4). In that case, the cost of no longer protecting established biomass was greater than the benefits of moving away from heatwaves. This suggests caution when relocating MPAs under climate-adaptive planning: if the goal is promoting resilience against heatwaves through protecting trophic cascades, moving MPAs to less-impacted areas is not always optimal. While placing MPAs in climate refugia may be better long-term, relocation could incur prohibitive short-term costs (including ecological and non-ecological; Green et al., 2014). If moving MPAs is a key objective, maintaining old MPAs until newer ones have rebuilt biomass may mitigate the short-term costs.

A key insight from our findings is that when establishing new MPAs, focusing protection on patches within heatwave-prone areas was the best strategy (for within-MPA effects), but when reconfiguring an established MPA network, protecting heatwave refugia was more effective. This difference arises because in the case of existing MPAs, there was a large difference in resilience inside versus outside MPAs (i.e., the median inside was 100%; Fig S3). When an old MPA was expanded in the non-impacted refugium habitat, existing kelp drift and spore spillover from that MPA patch was able to support its new MPA neighbor to dramatically increase resilience. This spillover, however, was less effective from an existing MPA in the heatwave-impacted habitat. By contrast, in the case of adding new MPAs where none previously existed, the time required to build up biomass (i.e., the transient dynamics) limited the capacity for resilience benefits. What effect existed was similar in magnitude in both the impacted and refuge parts of the coastline, but because the baseline scenario (no MPAs) had lower resilience in the impacted zone, the relative benefit of MPAs was greater there (Fig S2). In other words, when expanding MPAs, doing so in the non-impacted refuge area was more effective both in absolute terms (higher resilience within MPA boundaries) and relative to the baseline scenario. But when adding MPAs, there was no spatial difference in absolute terms, but adding MPAs to the heatwave-impacted area was better relative to the no-action baseline.

It is important to recognize that our results regarding MPA planning apply to a specific ecological context: MPAs that protect fishery-targeted grazer (i.e. urchin) predators to reduce the likelihood of heatwave-driven kelp forest collapse through overgrazing. The key elements driving our findings are that the urchin predator is fished (which is not the case in all kelp forests; Kumagai et al., 2024; Spiecker et al., 2025), that the dispersal of kelp spores and drift kelp is highly local, relative to the connectivity of fish and urchin larvae, and that heatwaves are the primary climate stressor. Heatwaves have emerged as a key threat to kelp forest resilience (Parnell et al., 2026), but there are other climate stressors in temperate systems that do not necessarily covary with temperature (Hamilton et al., 2023). 2023). As minimizing exposure to one stressor may increase exposure to another (Bruno et al., 2018), our findings need to be considered in a broader planning approach that balances MPA configurations to achieve clear goals.

The general tri-trophic framework we have developed could also apply to some coral reef systems where protecting a functionally diverse herbivore community supports coral persistence (Rasher et al., 2013). System-specificity aside, an important general lesson from our analysis is the value of an ecologically mechanistic modeling approach for climate-informed conservation planning. By explicitly modeling trophic dynamics and transient population dynamics, we revealed subtle and counterintuitive guidance for MPA placement (e.g., the trade-offs involved in expanding existing MPAs in refuges versus adding new MPAs to heatwave-impacted habitats) that are not possible from only examining distributions of forecasted physical ocean statistics (e.g., sea surface temperatures) and choosing to protect ‘refuge’ habitat (Green et al., 2014). As this study demonstrates, it is important to explicitly account for the spatiotemporal constraints of connectivity and transient population dynamics when considering climate-informed spatial management.

## Methods

### Three-species, multi-patch population model

Here we extend the three-species, single-patch model from Hopf et al. (2025) – which captures the interactions between giant kelp, herbivorous purple urchins, and a fishery-targeted urchin-eating predatory fish (California Sheephead) – to a 16-patch along-coast population model with connectivity between all patches. To account for seasonal variations in ecological processes (recruitment, kelp growth, urchin grazing, and heatwave effects), we use seasonal (3-month) time steps. The model captured the characteristic kelp forest dynamics in which local overgrazing by urchins leads to urchin barrens at the patch scale. Kelp spore recruitment and external drift supply can then rescue extinct patches under low urchin densities. Here we give an overview, but details are provided in the Supplemental Information, and code is available at DOI: [[TBA]].

At the patch level (∼1 km^2^), our model is comprised of two linked sub-models: (1) a stage-based kelp-urchin component tracking biomass of juvenile/adult kelp (standing and drift) and juvenile/adult urchins (exposed and hiding), and (2) a predator integral projection model (IPM) tracking Sheephead length-abundance converted to biomass. Our model incorporates behavioral switching where urchins passively consume drift kelp while hiding but emerge to graze standing kelp when drift becomes scarce, with exposure modeled as a declining function of drift availability (Randell, 2022; Rennick et al., 2022). We used a Type-II functional response for urchin grazing and a Type-I response for Sheephead predation on adult urchins, with higher predation mortality on exposed versus cryptic urchins (Nichols et al., 2015). When kelp was absent for >3 months, Sheephead ceased consuming nutritionally-depleted urchins (Liebergesell, 2022).

We modelled an effectively infinite linear coastline (the first patch neighbors the last patch), representing the Southern California coast, where kelp forest patches are interspersed with sandy habitat. Kelp forest patches were demographically connected through kelp spore and drift dispersal, and purple urchin and Sheephead larval dispersal. We used a 1km^2^ patch size, which encompasses adult Sheephead (Topping et al., 2006) and urchin (Rogers-Bennett & Okamoto, 2020) home ranges, and captures the majority (∼65%; Gaylord et al., 2002), but not all, of kelp spore dispersal. Kelp spore dispersal between patches followed a data-parameterized negative exponential function (Gaylord et al., 2002), and we assumed that kelp drift only dispersed between nearest-neighbor patches (Figurski, 2010; Hobday, 2000). Urchin and Sheephead larval dispersal were assumed to be well-mixed among patches, reflecting the long larval durations and distances of both species (Alonzo et al., 2004; Kinlan & Gaines, 2003). Stochasticity was captured through year-to-year variation in recruitment for all species, with recruitment each year drawn randomly from data-parameterized normal distributions. Where possible, the model is parameterized using published data from the Channel Islands, southern California, otherwise data from comparable regions are used (see SI for more details).

### Modelling heatwaves & MPA strategies

Reflecting the known effects of heatwaves on kelp and urchins, we implemented heatwaves as a period of reduced kelp recruitment (Hollarsmith et al., 2020) and growth (Zimmerman & Kremer, 1986), and increased urchin grazing rates (Spindel, 2023). As an illustrative scenario, we assumed that half of the modelled coastline experienced heatwaves, and that there was a 0.25 probability that the heatwave would occur each year, reflecting the expected century-scale increase in the frequency of heatwaves in the California Current (Shi et al., 2021).

To establish initial conditions, we ran the two modelled baseline scenarios (without MPAs and with 12.5% MPA coverage, a value chosen because it is approximately the coverage of MPAs in California waters) for 40 years without the increased heatwave effect. Given that we parameterized the model (not including the modelled heatwave effects) using data primarily collected before the 2014-2016 Californian heatwave, we implicitly assume that the initial conditions reflect baseline climate levels of heatwaves. We then ran the model for 20 years with the increased heatwave effects and intensity under a range of MPA scenarios (Fig 2), simulating 5,000 replicates of each scenario. In scenarios with increased in MPA coverage, Sheephead fishing pressure was evenly redistributed to the patches that remained open to fishing.

### Resilience calculations

We used the proportion of time kelp biomass was above a minimum threshold (1% of no-heatwave, kelp-forested state) during the simulation run (20 years) as our resilience response variable. We plotted the distributions of resilience at the patch and whole coast level (SI Figs) and focused on the change in median resilience between the baseline and alternative scenarios for both the with and without old MPA cases.

## Acknowledgements

This work was funded by the David and Lucile Packard Foundation (award 2022-74730), the California Ocean Protection Council (agreements C0874012 and C0874015), Oregon Department of Fish and Wildlife (contract IGA 251-23) and in part by the Oregon Agricultural Experiment Station with funding from the Hatch Act capacity funding program, award numbers NI25HFPXXXXXG022 and/or NI25HMFPXXXXG029, from the USDA National Institute of Food and Agriculture. This is publication XX of the Partnership for Interdisciplinary Study of Coastal Oceans, funded primarily by the David and Lucile Packard Foundation

## Footnotes

1 Note that we are referring to the ‘spillover’ of kelp spores and drift, rather than the traditional MPA spillover of fishery-targeted fish. In our model, adult fish remain within their recruited patch.

