## Supplemental Information for "Tradeoffs in planning marine protected areas for kelp forest resilience: protecting climate refugia is not always the best solution"

1. Supplemental Methods 3
1.1. Model overview 3
1.2. Sheephead sub-model 3
1.3. Urchin-kelp sub-model: purple urchins 7
1.4. Urchin-kelp sub-model: giant kelp 11
2. Supplementary Figures 19
3. References 23

### 1. Supplemental Methods

#### 1.1. Model overview

Here we extend the three-species, single-patch model from Hopf et al. (2025) to a 16-patch along-coast population model with connectivity among all patches. Note that these Supplemental Methods have been adapted from Hopf et al. (2025), with the authors' permission.

Our model captures the interactions between giant kelp (*Macrocystis pyrifera*; hereafter 'kelp'), herbivorous purple urchins (*Strongylocentrotus purpuratus*), and the fishery-targeted, urchin-predating Californian Sheephead (*Bodianus pulcher*). At the patch level, the model comprises two sub-models: 1) a predatory fish integral-projection sub-model (IPM; following Hopf & White, 2023), which tracks the abundance of Sheephead of all lengths over time; we then translate length-abundance into biomass using an allometric relationship; and 2) an urchin-kelp stage-based sub-model that tracks biomasses of key life-stages: juvenile and adult standing kelp, drift kelp (detached kelp fronds), juvenile urchins, and exposed and hiding adult urchins.

Patches were separated by sandy substrate along a linear, infinite coastline (such that patch 1 and patch 16 are neighbors). Patches were demographically connected through kelp spore and drift dispersal, and purple urchin and Sheephead larval dispersal. We assumed patches had an area of approximately 1 $km^2$, which encompasses adult Sheephead (Topping et al., 2006) and urchin (Rogers-Bennett & Okamoto, 2020) home ranges, and would capture the majority (~65%; Gaylord et al., 2002), but not all, of kelp spore dispersal, measuring from the center of the patch.

The model runs over seasonal (3-monthly) time steps, following the Northern Hemisphere seasons: winter (December – February), spring (March – May), summer (June – August), and autumn (September – November).

Where possible, the model is parameterized using data from the Channel Islands, southern California, otherwise data from comparable regions are used (see Tables S1.1 & S1.3 for specifics). Where parameters have been arbitrarily set to match trends or densities observed in the Channel Islands, this was done under the baseline 12.5% MPA coverage scenario. Data sources are either published literature (see Tables S1.1 & S1.3) or published datasets (PISCO; access through DataONE). Models were run in MATLAB (The MathWorks Inc., 2022). All code can be found at DOI: [[TBA]].

#### 1.2. Sheephead sub-model

To model Sheephead population dynamics over time, we used an integral-projection model (IPM; Ellner & Rees, 2006; Rees et al., 2014; White et al., 2016), which models dynamics in a size-structured, discrete-time manner. This model structure captures well changes in population size structures over time, which translates into changes in predation of urchins, as larger Sheephead consume more urchin biomass. Our model structure closely follows Hopf and White (2023). All model parameter descriptions and values for the Sheephead sub-model are in Table S1.1.

The Sheephead IPM tracks the state of patch *p* in terms of its size distribution, $n_p(x,t)$, which is the density function of individuals of size *x* in season *t*. The population density at time $t+1$ is the current population density multiplied by the probability density of moving from size *x* to size *y*, $K(y,x)$, plus new recruits, integrated over all biologically reasonable sizes ω:

$$n_p(y,t+1) = \int_\omega K(y,x)\, n(x,t)dx + R_S\, \rho(y)\, \theta_S.$$

The kernel $K(y,x)$ is a product of the growth kernel for survivors, $G(y,x)$ (a von Bertalanffy growth function), and survivorship, $P(y,x)$, which is size-dependent:

$$P(y,x) = e^{-[M-\phi(x)F]}G(y,x)\ ,$$

where *M* is instantaneous natural mortality rate per season, *F* is the fishing mortality rate per season, and $\phi(x)$ is the selectivity function describing the vulnerability of size *x* fish to harvest. The selectivity function $\phi(x)$ is zero below the legal size limit and one above it.

Assuming a closed population (at the entire-coast metapopulation scale) with well-mixed larval dispersal among patches, the total number of incoming recruits ($R_{S,t}$) to a given patch is a function of size-based-fecundity ($f(x)$), total adult biomass at given size (summed across the whole coastline), the yearly variability in incoming recruits ($\sigma_S$), and the recruitment timing function ($\theta_S$) for Sheephead, which captures autumn peaks in Sheephead recruitment (Alonzo et al., 2004; Cowen, 1991, 1990), all divided by the number of patches.

Arriving recruits compete within their cohort for resources, following a Beverton-Holt density-dependence function, where the per-capita survival of a recruit ($s_{SR}$) is

$$s_{SR} = \frac{\alpha_1}{1+\frac{\alpha_1 R_{S,t}}{\beta_1}}$$

To parameterize the Beverton–Holt function, we (1) set the slope at origin ($\alpha_1$) so that the population collapses if fishing decreases average lifetime egg production below 25% of the unfished maximum (similar to assumptions about stock-recruit relationships made for many U.S. Pacific coast fishes; Botsford et al., 2019) and (2) set the theoretical average maximum density of recruits ($\beta_1$) across patches (under the 12.5% baseline MPA scenario) to achieve average Sheephead densities observed in

Channel Islands kelp forests[1]. Finally, the total number of recruits that survive density dependence is distributed across the probability density function for initial recruit size $\rho(y)$.

The total biomass of Sheephead predating urchins in a given season *t*, in a given patch *p* ($N_{t,p}$) is therefore the product of the abundance density function, $n_p(y,t)$, and the weight-at-length function, $W(y)$, integrated for all sizes greater than then average size of Sheephead that are able to effectively consume urchins, $x_{pred}$:

$$N_{t,p} = \int_{x_{pred}}^{\infty} n(y,t)\, W(y)\, dy.$$

$W(y) = g\, y^h$, where *g* and *h* are functional shape parameters.

To remove the effects of transient dynamics in predator biomass densities, the first 40 years (long enough for transient dynamics to settle) of the Sheephead model were discarded. We assume that the fishery, and MPAs in the 12.5% MPA coverage baseline scenario, had been in place long enough that the fished population has reached a time-averaged equilibrium.

[1] See PISCO_kelp-urchin-Sheephead_data.Rmd file for details

**Table S1.1** Parameter descriptions, values, and references (where applicable) for the Sheephead sub-model.

| Symbol | Description | Value | Reference/Source |
|---|---|---|---|
| $M$ | Adult instantaneous natural mortality rate (per season) | 0.2/4 | Alonzo et al. (2004) |
| $F$ | Adult instantaneous mortality rate due to fishing (per season) | 0.2/4 | Alonzo et al. (2004) |
| $x_F$ | Legal size limit | 30 cm | Alonzo et al. (2004) |
| $\boldsymbol{x_{pred}}$ | Minimum fish size able to effectively consume urchins | 20 cm | Hamilton and Caselle (2015)<br>Selden et al. (2017a) |
| $\boldsymbol{f(x)}$ | Fecundity of fish size $x$<br>$f(x) = vx^w$<br>$v$ = constant shape parameter<br>$w$ = exponent shape parameter | <br><br>7.04<br>2.95 | Alonzo et al. (2004) |
| $\sigma_S$ | standard deviation of incoming recruits | 0.7188 | PISCO data* |
| $\boldsymbol{\beta_1}$ | Theoretical maximum density of recruits | 6*10^5 | Patch-average value set to achieve average unfished of Sheephead densities in Channel Islands kelp forests, based on PISCO data* |
| $\boldsymbol{\theta_S}$ | Sheephead recruitment timing function | [0 0 0.05 0.95]+ | Cowen (1990, 1991)<br>Alonzo et al. (2004) |
| $g$ | Weight-at-length shape parameter | $2.7 \times 10^{-5}$ | Alonzo et al. (2004) |
| $h$ | Weight-at-length shape parameter | 2.86 | Alonzo et al. (2004) |

* See PISCO_kelp-urchin-Sheephead_data.Rmd file for details.

+ Seasonal variations, in the order winter, spring, summer, autumn

*1.3. Urchin-kelp sub-model: purple urchins*

See Table S1.2 for state variables, and Table S1.3 for model parameters descriptions and values for the urchin-kelp sub-model.

As we are including a kelp-drift feeding switch in urchin behavior, we chose a stage-based matrix model for the kelp-urchin sub-model, which easily allows attribution of different demographic rates to different life-history or behavioral stages. We assumed a closed population (at the coast scale) for urchins, with well-mixed larval dispersal among patches. This is a fair assumption as urchins have long larval durations and distances (Arafeh-Dalmau et al., 2022; Kinlan & Gaines, 2003).

A vec-permutation approach (Caswell, 2001; Hunter & Caswell, 2005) suits well our multi-patch, stage-based matrix approach. Changes over time (seasons) in the urchin population can be written as

$$\mathbf{u}_{t+1} = \mathbf{P}^{\mathrm{T}}\mathbb{D}_{\mathrm{U}}\mathbf{P}\mathbb{M}_{\mathrm{U}}\,\mathbf{u}_{t}.$$

The urchin population vector, $\mathbf{u}$, includes the biomass densities of each urchin stage – juveniles (*J*), cryptic adults (*C*), and exposed adults (*E*), respectively – in each patch. Demographic and dispersal processes are captured in separate block diagonal matrices, which are combined through $\mathbf{P}$, the vec-permutation matrix. The block diagonals on $\mathbb{M}_{\mathrm{U}}$ (demography) and $\mathbb{D}_{\mathrm{U}}$ (dispersal) are 3 x 3 (3-stages) and 16 × 16 (16-patches) projection matrices for the demography of patch *p* ($\mathbf{M}_{\mathbf{U,t,p}}$) and dispersal of stage *i* ($\mathbf{D}_{\mathbf{U,t,i}}$), respectively.

Following the assumptions of well-mixed urchin larval dispersal and no adult movement between patches, the dispersal matrix for juveniles ($\mathbf{D}_{\mathbf{U,t,J}}$) is a 16 x 16 constant matrix with all elements equal to 1/16, and the dispersal matrices for cryptic and exposed adults ($\mathbf{D}_{\mathbf{U,t,C}}$ & $\mathbf{D}_{\mathbf{U,t,E}}$) are 16 x 16 identity matrices.

The urchin demographic transition matrix for patch *p* ($\mathbf{M}_{\mathbf{U,t,p}}$) is

$$\mathbf{M}_{\mathbf{U,t,p}} = \begin{matrix} \\ \mathit{juvenile} \\ \mathit{cryptic} \\ \mathit{exposed} \end{matrix} \begin{matrix} \begin{matrix} \mathit{juvenile} & \mathit{cryptic} & \mathit{exposed} \end{matrix} \\ \begin{bmatrix} (1-g_U)\,s_J & R_{U,t} & \psi\,R_{U,t} \\ g_U\,s_{C,p}\,(1-\phi_{\mathrm{t,p}}) & s_{C,p}\,(1-\phi_{\mathrm{t,p}}) & s_{C,p}\,(1-\phi_{\mathrm{t,p}}) \\ g_U\,s_{E,p}\,\phi_{\mathrm{t,p}} & s_{E,p}\,\phi_{\mathrm{t,p}} & s_{E,p}\,\phi_{\mathrm{t,p}} \end{bmatrix} \end{matrix},$$

where $R_{U,t}$ is the biomass of urchin recruits per kilogram of urchin adult, $g_U$ is the proportion of juvenile biomass maturing to adults in a season, $s_i$ is the proportion of stage *i* surviving the season, and $\phi_{\mathrm{t}}$ is the proportion of urchin biomass exposed in the open, grazing adult standing kelp in a given season *t* (the behavior switching function). We explain these parameters further in the following paragraphs. Finally, $\psi$, is a step function that turns off predation of urchins and urchin reproduction when a kelp barren state (defined as standing kelp biomass less than a minimum threshold, $k_{JA}^{min}$) has existed for great than 3 months. Starving barren-state urchins have significantly reduced gonad weights, leading to minimal reproductive capacity (Okamoto, 2014) and Sheephead actively avoid staving barren-state urchins (Eurich et al., 2014), which are nutritionally poor and adhere strongly to the substrate (Liebergesell, 2022).

Urchin recruitment varies seasonally, with ~90% of recruitment occurring between March to July (spring into summer; Okamoto et al., 2020a), and shows interannual variation (Ebert et al., 1994). The biomass of urchin recruits per kilogram of adults ($R_{U,t}$) is:

$$R_{U,t} = norm\left(\overline{R_U}, \sigma_{RU}\right) \theta_{U,t} \left(s_J\right)^{\tau},$$

where $\overline{R_U}$ is the yearly mean biomass of urchin recruits per kilogram of adult, $\sigma_{RU}$ is the total the yearly variance of incoming urchin recruits, and $\theta_{U,t}$ is the timing function for urchin recruitment (capturing seasonal variation and summing to one over the year). Note that $\overline{R_U}$ is arbitrary and we set this parameter to achieve average urchin densities observed in Channel Islands kelp forests[2]. Lastly, since planktonic larval durations (PLDs) for urchins are typically less than 3 months (40-90 days; Arafeh-Dalmau et al. 2022), recruits will operationally spend a portion of time as juveniles during a time step. We therefore also included a juvenile survival term ($s_J$) that is discounted by the proportion of time spent as a recruited juvenile ($\tau$): $\tau = 1 - \frac{PLD}{91}$. Note, that when $\tau = 1$, $\left(s_J\right)^{\tau} = s_J$.

Growth and maturation is calculated as

$$g_U = \frac{1}{(4 * g_{mat})},$$

reflecting the probability that a juvenile will mature that season given the average age of urchin maturation ($g_{mat}$).

Urchin survival ($s_i$) is a function of natural survival rates ($M_i$) for juveniles and adults, and the rate of predation per kg of local predators ($P_i$) for adults, such that

$$s_J = e^{-M_J}$$

$$s_{C,p} = e^{-M_H - P_C N_{t,p}}, \text{ and}$$

$$s_{E,p} = e^{-M_E - \psi P_E N_{t,p}}, \psi = \begin{cases} 0, & \sum_{T=1}^{2}\left(k_{J,t-T} + k_{A,t-T}\right) < k_{JA}^{min} \\ 1 \end{cases}.$$

Outputs from the Sheephead predator sub-model provided the input values for $N_{t,p}$. Here we are assuming linear (Type-I) predation of urchins, where per kg predation by Sheephead is constant. This reasonable for Sheephead, as urchins only make up a small proportion of Sheephead diets and they would be unlikely to reach high, saturating feeding rates (Cowen, 1983). However a Type-II or Type-III functional response may be more appropriate for other predators such as rock lobsters (Dunn &

[2] See PISCO_kelp-urchin-Sheephead_data.Rmd file for details

Hovel, 2020). We also assume that Sheephead do not predate juvenile urchins as juveniles are typically afforded protection by hiding in the spines of adults (Tegner & Dayton, 1981).

The urchin behavioral feeding switch function ($\phi_{t,p}$) captures the shift between urchins cryptically grazing on drift kelp when resources are plentiful, to leaving rocky crevices and targeting standing kelp when drift resources are relatively low (Ebeling et al., 1985; Kriegisch et al., 2019; Randell, 2022; Rennick et al., 2022a). This behavior will vary by patch. Using a declining logistic function to model the proportion of urchins grazing upon kelp as a function of drift density, Randell (2022) demonstrated how this behavioral switching can drive alternative kelp-forested and urchin-barren states. We use the same functional form as Randell (2022) but normalize drift density to the total consumptive capacity of urchins (total grazing capacity of the urchin biomass; Rennick et al., 2022a) in patch *p*, a given season. Consequently, the inflection point beyond which more urchins graze standing kelp than drift is when the consumptive capacity exceeds drift supply (i.e., grazing > drift; Figure S1.1). In doing this, we allow for the influence of changing grazing rates with marine heatwaves on urchin switching behavior. A such, the proportion of urchins exposed, grazing standing kelp ($\phi_{t,p}$) at time *t* is

$$\phi_{t,p} = e^{-v_1\left(\frac{\overbrace{k_{D,t}}^{\text{drift kelp density}}}{\underbrace{b_{D,C}\,(u_{C,t,p}+u_{E,t,p})}_{\text{urchin consumptive capacity}}}\right)^{v_2}},$$

where the following are satisfied:

$$v_1 = \frac{v_2-1}{v_2} \quad \text{and} \quad (v_2-1)e^{\frac{1-v_2}{v_2}} = w,$$

$w$ is the slope around the inflection point, and $b_{D,C}$ is the maximum grazing rates of drift kelp by cryptic urchins (see below).

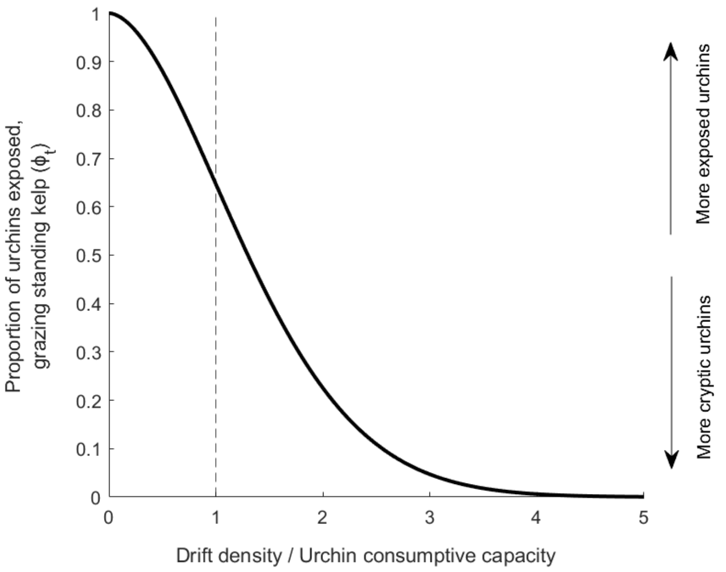


**Figure S1.1** urchin behavioral feeding switch function ($\phi_t$). As the availability of drift kelp to urchin consumptive capacity (biomass of urchins x urchin grazing rate) decreases (left-to-right along the x-axis) more urchins shift to grazing standing kelp. The rate of change of this shift is determined by the slope of the curve ($w_2$), here $w_2 = 0.5$.

##### *1.4. Urchin-kelp sub-model: giant kelp*

Following the same multi-patch, stage-based matrix approach, the population vector for kelp biomass at time *t* ($\mathbf{k}_{t+1}$) is

$$\mathbf{k}_{t+1} = \mathbf{P}^{\mathrm{T}} \mathbb{D}_{\mathrm{K}} \mathbf{P} \mathbb{M}_{\mathrm{K}} \, \mathbf{k}_{t+1} \, ,$$

where the $\mathbb{D}_{\mathrm{K}}$ is the 16 x 16 kelp dispersal block diagonal matrix, and $\mathbb{M}_{\mathrm{K}}$ is the 3 x 3 kelp demography block diagonal. The three kelp life-stages are standing juvenile sporophytes (*J*), standing adult sporophytes (*A*), and detached kelp fronds ('drift kelp'; *D*). We define juvenile kelp as sporophytes that are visible to the naked eye, but not mature (0-3 months old; Schiel & Foster, 2015). Juveniles are considered standing kelp and are, therefore, grazed by urchins and contribute to the production of drift.

To quantify kelp spore dispersal between patches, we fitted an negative exponential curve to published data on kelp spore dispersal (Gaylord et al., 2002)[3], and calculated the probability of travelling the discrete distances among patches. Propagules that landed on the sandy substrate between patches were considered lost from the system, and the infinite coastline meant that, for example, propagules from patch 1 could reach patches 16, 15, 14, etc. As such, the dispersal matrix for kelp juveniles ($\mathbf{D}_{\mathbf{K,t,J}}$) had decreasing probabilities away from the main diagonal, with high self-retention within a patch.

Given that adult kelp stipes are fixed to the substrate, the dispersal matrix for kelp adults ($\mathbf{D}_{\mathbf{K,t,A}}$) is a 16 x 16 identity matrix. The dispersal of kelp drift is highly variable in space and time (Figurski, 2010; Hobday, 2000). In this along-coast model, we assume that drift that is not retained within a patch is either lost off-shore, washed onto beaches, or disperses to the nearest neighboring patch. Therefore, dispersal matrix for kelp drift ($\mathbf{D}_{\mathbf{K,t,D}}$) is a 16 x 16 matrix:

$$\mathbf{D}_{\mathbf{K,t,D}} = \begin{bmatrix} d & \frac{1-d}{2} & 0 & 0 & \cdots & 0 & 0 & 0 & \frac{1-d}{2} \\ \frac{1-d}{2} & d & \frac{1-d}{2} & 0 & \cdots & 0 & 0 & 0 & 0 \\ 0 & \frac{1-r_D}{2} & d & \frac{1-d}{2} & \cdots & 0 & 0 & 0 & 0 \\ 0 & 0 & \frac{1-d}{2} & d & \cdots & 0 & 0 & 0 & 0 \\ \vdots & \vdots & \vdots & \vdots & \ddots & \vdots & \vdots & \vdots & \vdots \\ 0 & 0 & 0 & 0 & \cdots & d & \frac{1-d}{2} & 0 & 0 \\ 0 & 0 & 0 & 0 & \cdots & \frac{1-d}{2} & d & \frac{1-d}{2} & 0 \\ 0 & 0 & 0 & 0 & \cdots & 0 & \frac{1-d}{2} & d & \frac{1-d}{2} \\ \frac{1-d}{2} & 0 & 0 & 0 & \cdots & 0 & 0 & \frac{1-d}{2} & d \end{bmatrix},$$

---

[3] See Gaylord_macro_dispersal.rmd for fitting

where $d$ is the proportion of drift biomass locally retained within a patch. Note that offshore and on-beach losses are included in the kelp demography matrix, below.

The demographic transition matrix ($\mathbf{M_{K,t,p}}$) in timestep $t$ and patch $p$ for kelp is

$$\mathbf{M_{K,t,p}} = \begin{array}{c} \\ juvenile \\ adult \\ drift \end{array} \begin{array}{c} \begin{array}{ccc} juvenile & adult & drift \end{array} \\ \begin{bmatrix} 0 & R_K\, \theta_{K,t}\, s_{Y,t,p}\, \gamma\, r_S\, \kappa_{Y,E,t,p} & 0 \\ g_K\, \gamma\, r_S\, \kappa_{A,E,t,p}\, (1-c) & g_K\, \gamma\, r_S\, \kappa_{A,E,t,p}\, (1-c) & 0 \\ c\, \kappa_{D,H,t,p} & c\, \kappa_{D,H,t,p} & r_D\, \kappa_{D,H,t,p} \end{bmatrix} \end{array}.$$

We explain all parameters in the following paragraphs (see also Table S1.3).

The biomass density of settled spores per kilogram of adults ($R_K$) includes zoospore production, survival and development through the gametophyte phase, successful fertilization, and settlement survival. To account for interannual variation in settling spores:

$$R_K = norm(\overline{R_K}, \sigma_{RK}),$$

where $\overline{R_K}$ is yearly mean biomass (per kilogram of adults) and $\sigma_{RK}$ is the yearly variance of incoming kelp spores.

In southern California, peaks in recruited juvenile sporophytes occur summer to autumn (July-November; Reed et al., 2008a), following winter peaks in spore production (Reed et al., 1996). Capturing this seasonal variation, $\theta_{K,t}$ (the kelp recruitment timing function) distributes recruitment proportionally over the seasons and sums to one each year.

Incoming kelp recruits to a patch compete for space and are negatively affected by shading of older standing adult kelp biomass (Nisbet & Bence, 1989; Reed, 1990; Stewart et al., 2009). As such, we used a combined Ricker and Beverton-Holt function, which captures both inter- and intra-cohort density dependent effects (White, 2009). Per biomass recruit survival at time $t$ in patch $p$ ($s_{Y,t,p}$) is, therefore,

$$\bar{s}_{Y,t,p} = \frac{(D-1)\, k_{A,t,p}\, e^{\beta D k_{A,t,p}}}{D\, Y_t\, e^{\beta D k_{A,t,p}} + e^{\beta k_{A,t,p}}\, ((D-1)\, k_{A,t,p} - D\, Y_t)},$$

where $\beta$ is the strength of density dependence, $D$ is the relative strength of inter- versus intra-cohort density-dependent processes ($0 < D < 1$), and $k_{A,t,p}$ is the biomass of standing adult kelp at time $t$ in patch $p$. Since $\beta$ is not quantifiable, we set this parameter to achieve average kelp densities observed in Channel Islands kelp forests[4]. Kelp-patch dynamics are not overly sensitive to the value of $D$ (Hopf

[4] See PISCO_kelp-urchin-Sheephead_data.Rmd file for details

et al., 2025), and we set $D$ relatively low ($D = 0.01$) reflecting the case where the effect of a kilogram of adults on recruit survival is far greater than that of a kilogram of recruits.

Since kelp dynamics occur at a smaller scale than our 1 ha model domain (i.e., at the local transect scale), we approximated the model landscape scale density dependent survival (following White, 2011) as

$$s_{Y,t,p} \approx \bar{s}_{Y,t,p} + 0.5\,\bar{s}''_{Y,t,p} + var\left(k_{A,t,p}\right)$$

where $var\left(k_{A,t,p}\right)$ is the spatial variance in the density of adult standing kelp in the local patch.

Reflecting fast maturation rates (on the order of months; Schiel & Foster, 2015), we assumed that juvenile sporophytes become adults, or contribute to drift, in the next season. Adult sporophyte growth over time is captured through $g_K$, the seasonal increase in standing kelp biomass, and $\gamma$, the relative change in standing (juvenile and adult) kelp biomass over the seasons.

Non-grazing loss of standing kelp biomass is captured in two terms: $c$, proportion of juvenile and adult standing kelp biomass converted to drift kelp in a time step, and $r_S$, the proportion of non-grazed standing kelp biomass that survives the season. $r_S$ accounts for storm and wave damage, senescing blade material, boat propeller damage, and dissolved material released from blades and stipes (Rassweiler et al., 2021). Drift kelp biomass that is neither retained nor dispersed to neighbouring patches (see $d$ above) is either lost through decomposition or removed from the system via processes such as transportation through wave action or storm events, where $r_D$ is proportion of drift biomass that remains in the population.

We assumed Type-II grazing functional response for standing and drift kelp by urchins, following Randell (2022). This is reasonable given that, like most herbivores following a Type-II saturating function, urchins require handling time between kelps stipes (Boada et al., 2017). $\kappa_{i,j,t,p}$ is the proportion of kelp stage $i$ (juvenile, adult, or drift), grazed by urchins stage $j$ (juvenile, cryptic adult, exposed adult), at time $t$, in patch $p$ such that:

$$\kappa_{i,j,t,p} = e^{-\Omega_{i,j,t,p}},$$

where the instantaneous rate of grazing $-\Omega_{i,j,t,p}$ of kelp stage $i$, by urchins stage $j$, at time $t$, in patch $p$, is a per-kilogram Type-II response :

$$\Omega_{i,j,t,p} = \frac{a_{i,j}}{1+\frac{1}{b_{i,j,t}}a_{i,j}\,k_{i,t,p}}\,u_{j,t,p},\ \ a_{i,t} > 0\ and\ b_i > 0.$$

$a_{i,j}$ is the per kilogram kelp mortality due to grazing by a kilogram of urchins in the absence of conspecifics (search/attack rate), and $b_{i,j,t}$ is the asymptotic (max) biomass of prey consumed by an urchin in a time step (1/handling and ingestion time) at time $t$ (this value changes under heatwave conditions). Note that exposed urchins eat recruits and adult standing kelp, and cryptic urchins only eat drift kelp.

**Table S1.2** State variables for the urchin-kelp sub-model. Each state variable is the biomass density (per 1km$^2$) at season $t$, in patch $p$.

| | | |
|---|---|---|
| *Urchins* | $u_{J,t,p}$ | Juvenile (small, sexually immature) urchins |
| | $u_{C,t,p}$ | Cryptic adult urchins |
| | $u_{E,t,p}$ | Exposed adult urchins |
| *Kelp* | $k_{J,t,p}$ | Juvenile standing kelp |
| | $k_{A,t,p}$ | Adult standing kelp |
| | $k_{D,t,p}$ | Drift kelp |

**Table S1.3** Parameter symbols, descriptions, values and references (where applicable) for the urchin-kelp sub-model. Values in red indicate values used during heatwave periods, and their associated references, where applicable. Unless otherwise specified, urchin data is for purple urchins (*Strongylocentrotus purpuratus*) and kelp data is for giant kelp (*Macrocystis pyrifera*; hereafter 'kelp'). Note that all relevant values are per hectare ($ha^{-1}$). Conversions used for urchin abundance to wet biomass were: 1) for size to biomass, Biomass (kg) = 0.0499 x test dimeter (cm) + 0.0019 (Ling et al. 2015), and 2) for number to biomass, Average purple urchin weight = 0.21kg (PISCO data*). Conversions used for kelp stipe numbers to wet biomass were: 1) for number to dry biomass, dry weight = 0.33stipes + 0.16 ((Reed et al., 2009)), and 2) from dry to wet biomass, wet-to-dry ratio = 0.094 ((Rassweiler et al., 2018)).

| | Symbol | Description | Baseline value ***(Heatwave value)*** | Units | References | Notes |
|---|---|---|---|---|---|---|
| **Urchins** | $\overline{R_U}$ | Mean biomass of settling recruits produced per kg of adults. This includes larval production (fecundity), dispersal and settlement survival. | 1.5 | $kg.yr^{-1}$ | | Value set to achieve a range of urchin biomasses that fall within observed values for Channel Islands kelp forests, based on PISCO data*. |
| | $\sigma_{RU}$ | Variance of incoming recruits | 0.621 | | | Normalised variance from PISCO data* for central and northern California. No urchin recruit data exists for Channel Islands. |
| | $\theta_{U,t}$ | Urchin recruitment timing function | [0.05 0.54 0.36 0.05]⁺ | | Okamoto et al. (2020b) | |
| | $g_{mat}$ | Average urchin maturation age | 2 | yrs | Gonor (1972) | |
| | $s_i$ | Proportion of urchin biomass stage *i* surviving the season | | | | See main text for relevant functions |
| | $M_j$ | Instantaneous natural mortality rate of stage *j* (not including predation) | $\begin{matrix} juvs \\ cryptic \\ exposed \end{matrix} \begin{bmatrix} 0.1 \\ 0.1 \\ 0.1 \end{bmatrix}$ | $season^{-1}$ | Russell (1987), Dunn et al. (2017) | |
| | $P_i$ | Instantaneous predation mortality rate of stage *j* (per kg predator) | $\begin{matrix} cryptic \\ exposed \end{matrix} \begin{bmatrix} 0.0065 \\ 0.013 \end{bmatrix}$ | $kg\ pred^{-1}.season^{-1}$ | Nichols et al. (2015), Selden et al. (2017b) | |
| | PLD | Urchin planktonic larval duration | 65 | days | Arafeh-Dalmau et al. (2022) | |
| | $w$ | Slope around inflection point of behaviour switching function | 0.5 | | | Value set so that kelp forest switches to urchin barren around observed urchin biomass thresholds (Ling et al., 2015). |
| | $\psi$ | Step function turning off predation of urchins in a kelp barren state | | | | See main text for relevant functions |
| | $k_{J+A}^{min}$ | Standing kelp density threshold below which a kelp barren state is declared | 1170 | kg | | Set to 1% of the average standing kelp densities observed at a persistent kelp forest location (Anacapa East) in the Channel Islands (~700kg.60m2/0.006 = ~ $1.17*10^5$kg.ha), based on PISCO data*. |
| **Kelp** | $\overline{R_K}$ | Yearly mean zoospore production, successful fertilization and | $4x10^4$ ***(baseline × ½)*** | $kg.year^{-1}$ | Dayton et al. (1984), Schiel and Foster | Value calculated as follows: |

| | Symbol | Description | Baseline value *(Heatwave value)* | Units | References | Notes |
|---|---|---|---|---|---|---|
| | | settlement of spore, *per kg adult kelp* | | | (2015), Hollarsmith et al. (2020) | • ~$10^{11}$ zoospores per plant (Schiel and Foster 2015, pg. 31)<br>• Halve value due to 50:50 sex ratio (Graham, 2007, pg. 47)<br>• ~1% survival probability to settled microsporophytes (Schiel and Foster 2015, pg. 77)<br>• ~20% survival probability for settled spores lasting 3 months (Dayton et al., 1984).<br>∴ $10^{11}$ x 0.5 x 0.01 x 0.2 = ~$10^8$ spores per plant<br>• We assumed that one recruit weighs, on average, 1g<br>• One adult plant = ~10kg wet (Rassweiler et al., 2018; Reed et al., 2009).<br>∴ ~$10^8$ spores per stipe = $10^8$ x 0.001 x 0.1<br>= $10^4$ kg recruits per kg adult (*per season*)<br>*Note that Schiel & Foster (2015, Fig 4.2A inset) have initial settlement rates on the order of $10^6$-$10^8$ spores.m$^2$, which converts to $10^2$-$10^4$ kg of spores per kg adult stipes.* |
| | $\sigma_{RK}$ | Variance of incoming settlers | 0.389 | | | Normalised variance from PISCO data* for Channel Islands California. |
| | $\theta_{K,t}$ | Kelp recruitment timing function | [0.1, 0.1, 0.4, 0.4]$^+$ | | Reed et al. (2008b) | |
| | $\beta$ | Strength of density dependence effect of shading by adults | 4x$10^{-6}$ | | | Value set to achieve a range of standing kelp biomasses that fall within observed values for Channel Islands kelp forests, based on PISCO data*. |
| | $D$ | Relative strength of inter- versus intra-cohort density-dependent processes | 0.01 | | White (2009) | |
| | $var(\overline{k_{A,t,p}})$ | Variance in the density of adult kelp in the local patch | 189,090 | | | Spatial variance at transect level, averaged over years, based on PISCO* kelp surveys at the Channel Islands. |
| | $g_K$ | Average increase in adult kelp biomass as it ages | 6.825 | kg.season$^{-1}$ | Stewart et al. (2009) | Conversion from average daily growth rate to seasonal (91 days): 0.075 x 91 = 6.825kg fronds per season |
| | $\gamma$ | Relative change in standing kelp biomass over the seasons | [1, 1, 0.8, 0.9]$^+$<br>***([1, 1, 0.5, 0.5])*** | | Zimmerman and Kremer (1986) | Based on average, pre-heatwave water temperatures observed in the Channel Islands. |
| | $r_S$ | Proportion of non-grazed standing kelp biomass remaining at the end of the season | 0.5688 | season$^{-1}$ | Rassweiler et al. (2018) | Conversion from average daily plant loss rate to proportion surviving a season (91 days): exp(-0.0062*91) = 0.5688. We assumed that biomass is proportional to number of plants. |

| | Symbol | Description | Baseline value *(Heatwave value)* | Units | References | Notes |
|---|---|---|---|---|---|---|
| | $c$ | Proportion of standing kelp biomass converting to drift (*drift production*) | 0.9 | season$^{-1}$ | Rassweiler et al. (2018), Rennick et al. (2022b) | Conversion from daily to seasonal proportion of biomass converted to drift = $1-(1-0.025)^{91}$. |
| | $d$ | Proportion of drift biomass retained locally, within a patch. ($1-d$) disperses to neighbouring patches | 0.7 | season$^{-1}$ | Hobday (2000), Figurski (2010) | ** Captured in the kelp dispersal matrix |
| | $r_D$ | Proportion of drift kelp that is not lost to decomposition or offshore/on-beach transportation. | 0.8 | season$^{-1}$ | Figurski (2010) | |
| | $a_{i,j}$ | Per kg mortality of kelp stage *i* due to grazing by an urchin, stage *j*, in the absence of conspecifics (*search/attack rate*) | $\begin{array}{r} \\ juv\ k \\ adult \\ drift \end{array} \begin{array}{c} hiding \quad exposed \\ \begin{bmatrix} 0 & 0.5 \\ 0 & 0.5 \\ 1 & 0 \end{bmatrix} \end{array}$ | kg kelp .kg urchin$^{-1}$ .season$^{-1}$ | Kriegisch et al. (2019) | No data is available to parameterise this, so we assumed very high attack rates with near-zero standing or drift densities. Kriegisch et al. (2019) findings suggest that attack rates for standing are less than that for drift as kelp has lower consumption than drift at high resource density (kelp forests). |
| | $b_{i,j,t}$ | Max kg of kelp stage *i* consumed by an urchin, stage *j*, in a time step (1/*handling time*) | $\begin{array}{r} \\ juv\ k \\ adult \\ drift \end{array} \begin{array}{c} hiding \quad exposed \\ \begin{bmatrix} 0 & b_t \\ 0 & b_t \\ b_t & 0 \end{bmatrix} \end{array}$<br><br>$b_t = 2.985 \times [1\ 0.9\ 1.15\ 1.2]^{+}$<br>$(\boldsymbol{b_t = 2.985 \times [1.15\ 1.05\ 1.2\ 1.3]})$ | kg kelp .kg urchin$^{-1}$ .season$^{-1}$ | Foster et al. (2015), Rennick et al. (2022b) Spindel (2023) | Kriegisch et al. (2019) findings suggest that the maximum feeding rate is the same for drift and standing kelp, as consumption rates were the same for drift and kelp in urchin barrens (when resources are low). We assume they are the same.<br>Experimental data (Randell, 2022; Rennick et al., 2022b) have a range of maximum feeding rate values and we used the average of these to obtain the baseline value.<br>• Rennick et al. (2022): 0.02 g detritus.g urchins$^{-1}$.day$^{-1}$.<br>Conversion = 0.02*1000/1000*91<br>= 1.82 kg kelp.kg urchin$^{-1}$.season$^{-1}$.<br>Urchins were taken from a kelp forest and fed for a week prior to experiments. Experiments were run at 14.5°C water temperature.<br>• Foster et al (2015): 1.14 g kelp.urchin$^{-1}$.day$^{-1}$.<br>Conversion = 1.14/1000*91<br>= 0.104 kg kelp.urchin$^{-1}$.season$^{-1}$<br>= 0.104/0.025<br>(mean experimental urchin weighed 25g)<br>= 4.15 kg kelp.kg urchin$^{-1}$.season$^{-1}$<br>Urchins were taken from an established urchin barren and starved for a week prior to experiments. Experiments were run at 11.6 to 16.3°C water temperature. |

| | Symbol | Description | Baseline value *(Heatwave value)* | Units | References | Notes |
|---|---|---|---|---|---|---|
| | | | | | | Finally, we used temperature experiments from Spindel (2023) to estimate relative change in max feeding rates over the seasons (based on average, pre-heatwave water temperatures observed in the Channel Islands.). |

[*] See PISCO_kelp-urchin-Sheephead_data.Rmd file for details. [+] Seasonal variations, in the order winter, spring, summer, autum

#### 2. Supplementary Figures

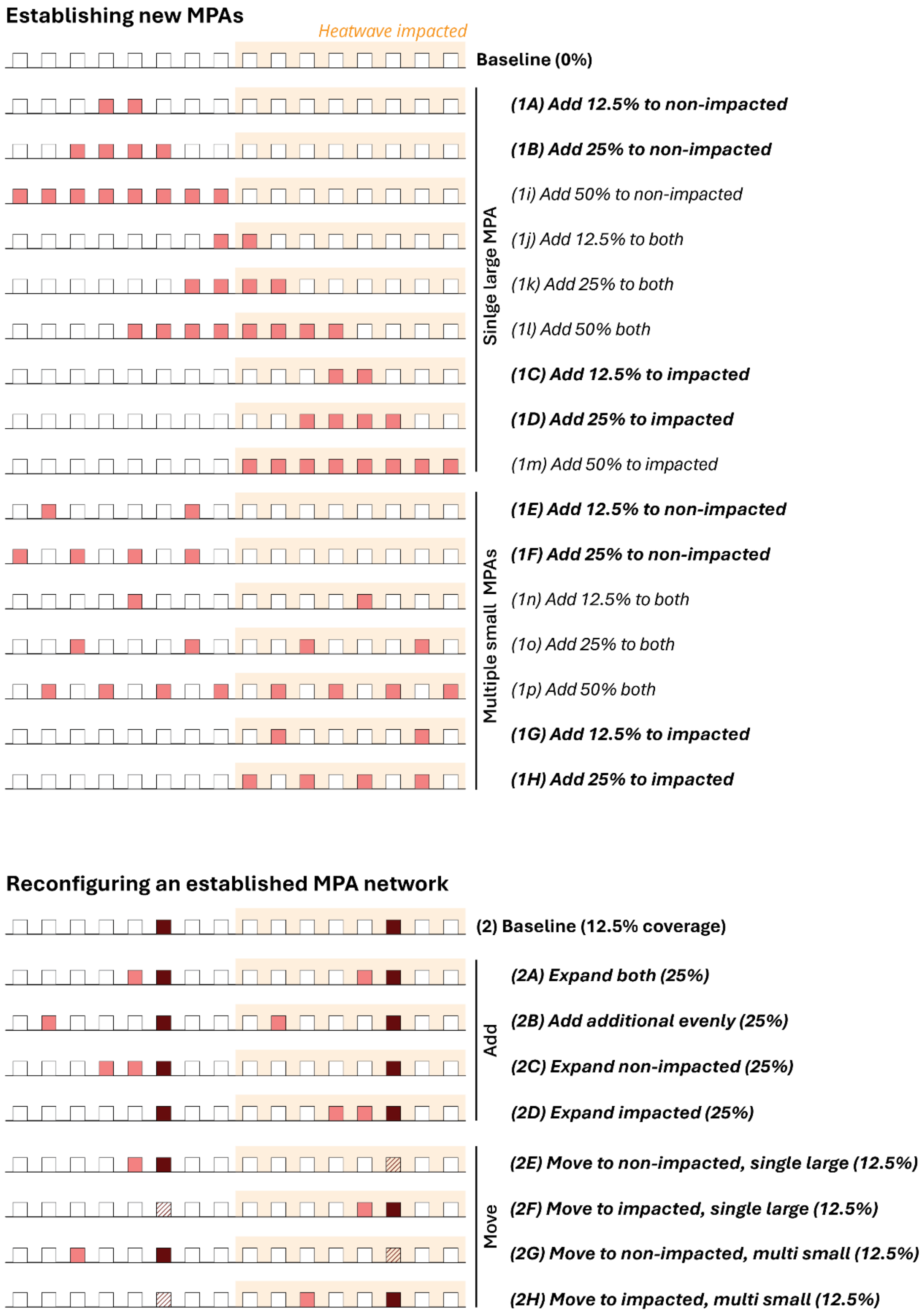


**Figure S1** Schematic of the modelled, linear coastline with all MPA scenarios considered in this study. Each square represents a single kelp forest patch covering 6.25% of the total kelp forest area, where light red squares are new MPAs, dark red squares are established MPAs, and white squares are fished patches. Orange regions indicate patches impacted by marine heatwaves. Scenarios with **bold text** are those presented in the main text.

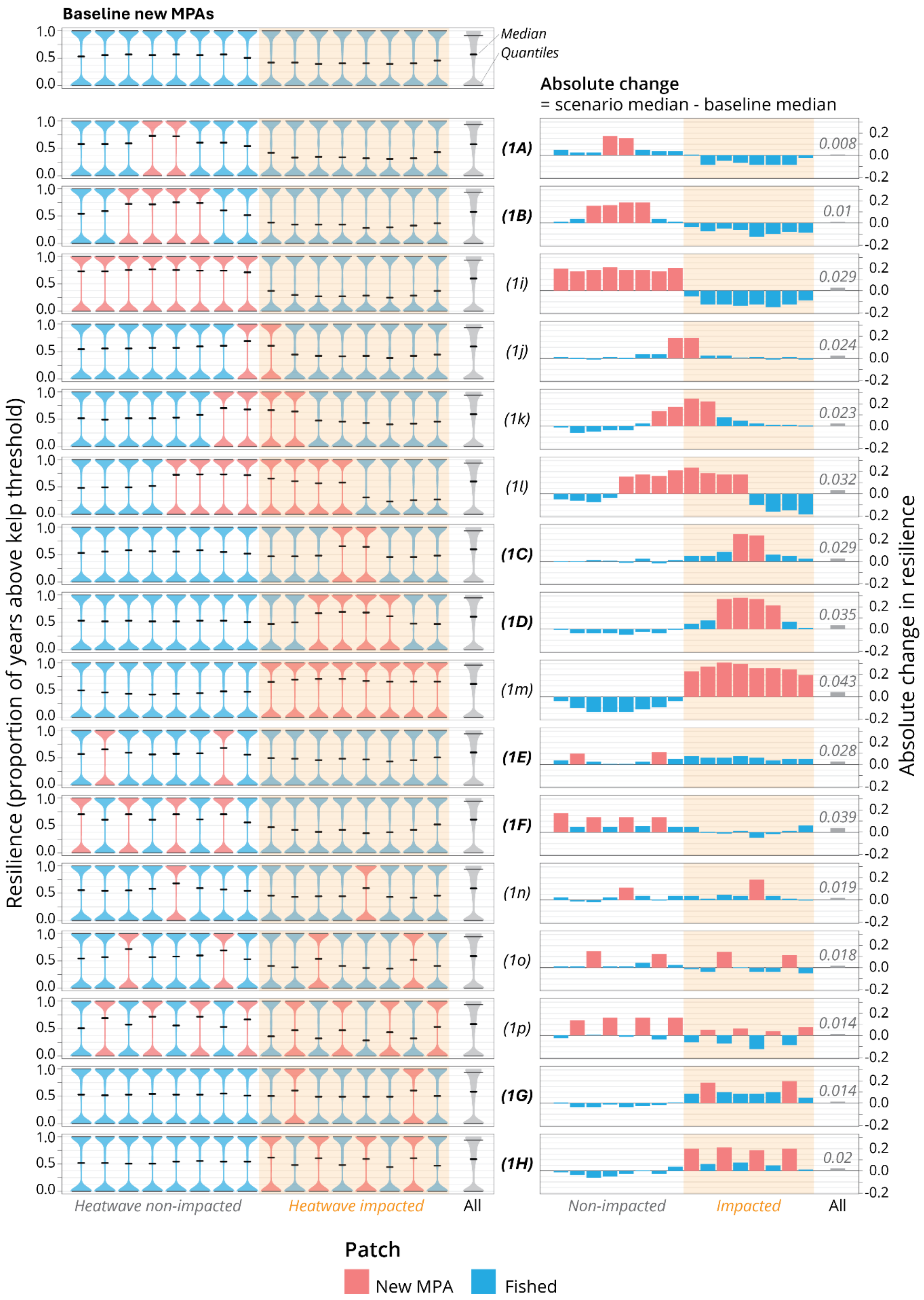
Baseline new MPAs
Median
Quantiles
Absolute change
= scenario median - baseline median
Resilience (proportion of years above kelp threshold)
Absolute change in resilience
(1A) 0.008
(1B) 0.01
(1i) 0.029
(1j) 0.024
(1k) 0.023
(1l) 0.032
(1C) 0.029
(1D) 0.035
(1m) 0.043
(1E) 0.028
(1F) 0.039
(1n) 0.019
(1o) 0.018
(1p) 0.014
(1G) 0.014
(1H) 0.02
Heatwave non-impacted
Heatwave impacted
All
Non-impacted
Impacted
All
Patch
New MPA
Fished

▲**Figure S2** Resilience effects when establishing new MPAs for all scenarios considered in this study. Median resilience (proportion of years kelp-forest patches spent above the kelp biomass threshold, which is 1% of the no-heatwave, kelp-forested state; left-hand column) and absolute change in the median resilience (right-hand column), 20 years post-MPA implementation, for the baseline (no MPAs) and alternative MPA scenarios. Each blue/red violin plot or bar indicates an individual patch, ordered along an infinite modelled coastline that is partially affected by marine heatwaves (orange box indicates impacted patches). Grey violin plots and bars are population means. Note that each patch covers 6.25% of the area. Letters relate to scenarios outlined in the top half of Figure S1, and scenarios with **bold text** are those presented in the main text.

▼**Figure S3** Resilience effects of reconfiguring an established MPA network. Median resilience (proportion of years kelp-forest patches spent above the kelp biomass threshold, which is 1% of the no-heatwave, kelp-forested state; left-hand column) and absolute change in the median resilience (right-hand column), 20 years post-MPA implementation, for the baseline (no MPAs) and alternative MPA scenarios. Each blue/red violin plot or bar indicates an individual patch, ordered along an infinite modelled coastline that is partially affected by marine heatwaves (orange box indicates impacted patches). Grey violin plots and bars are population means. Note that each patch covers 6.25% of the area. Letters relate to scenarios outlined in the top half of Figure S1.

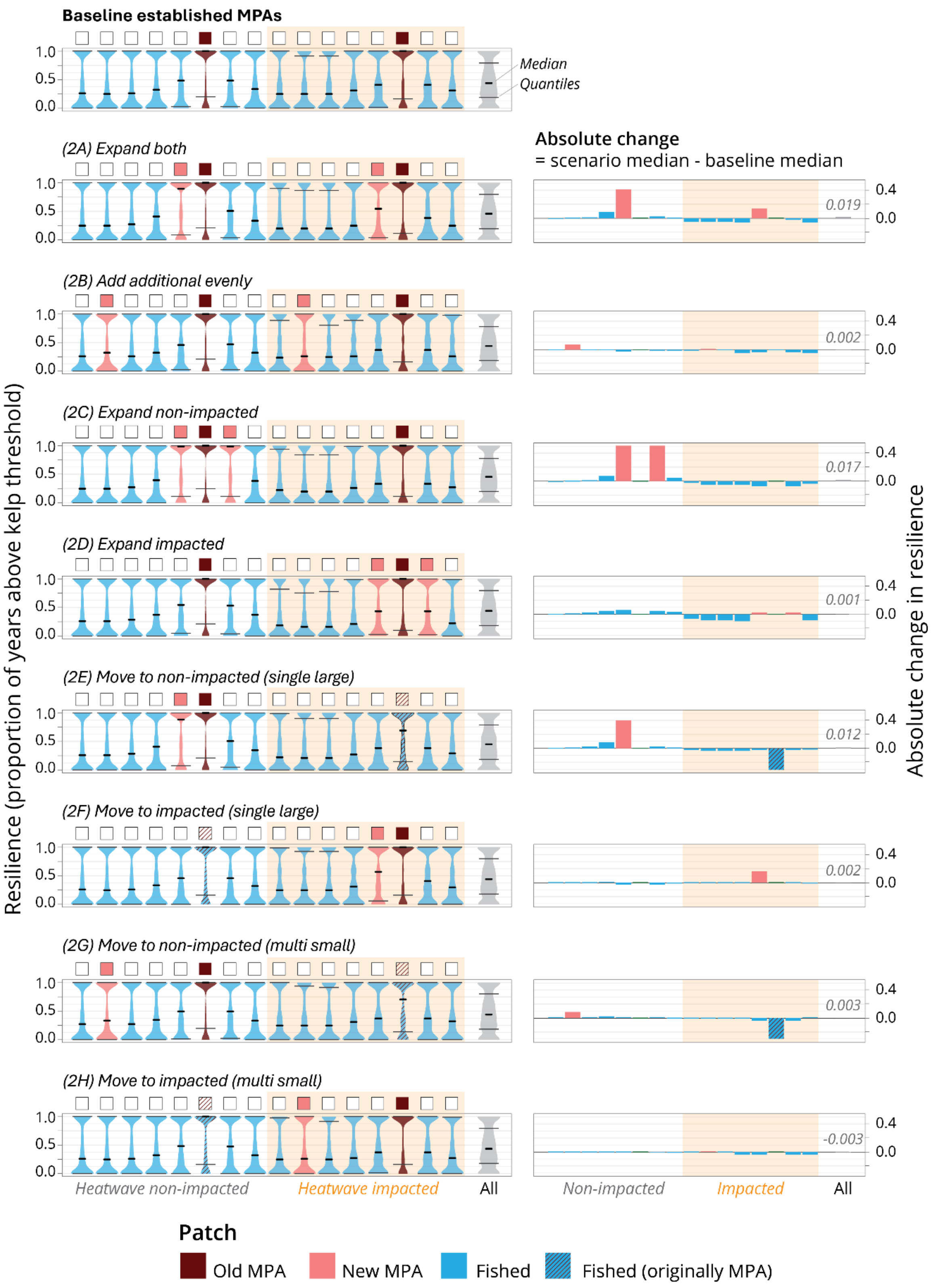
Baseline established MPAs
Median
Quantiles
(2A) Expand both
(2B) Add additional evenly
(2C) Expand non-impacted
(2D) Expand impacted
(2E) Move to non-impacted (single large)
(2F) Move to impacted (single large)
(2G) Move to non-impacted (multi small)
(2H) Move to impacted (multi small)
Resilience (proportion of years above kelp threshold)
1.0
0.5
0.0
Heatwave non-impacted
Heatwave impacted
All
Absolute change
= scenario median - baseline median
0.4
0.0
0.019
0.002
0.017
0.001
0.012
0.002
0.003
-0.003
Absolute change in resilience
Non-impacted
Impacted
All
Patch
Old MPA
New MPA
Fished
Fished (originally MPA)
